# The global root exudate carbon flux

**DOI:** 10.1101/2024.02.01.578470

**Authors:** Nikhil R. Chari, Shersingh Joseph Tumber-Dávila, Richard P. Phillips, Taryn L. Bauerle, Melanie Brunn, Benjamin D. Hafner, Tamir Klein, Sophie Obersteiner, Michaela K. Reay, Sami Ullah, Benton N. Taylor

## Abstract

Root exudation, the export of low-molecular weight organic carbon (C) from living plant roots to soil, influences microbial activity, nutrient availability, and ecosystem feedbacks to climate change, but the magnitude of this C flux at ecosystem and global scales is largely unknown. Here, we synthesize *in situ* measurements of root exudation rates and couple those to estimates of fine root biomass to estimate global and biome-level root exudate C fluxes. We estimate a global root exudate flux of 15.2 PgC y^-1^, or about 10% of global annual gross primary productivity. We found no differences in root mass-specific exudation rates among biomes, though total exudate fluxes are estimated to be greatest in grasslands owing to their high density of absorptive root biomass. Our synthesis highlights the global importance of root exudates in the terrestrial C cycle and identifies regions where more *in situ* measurements are needed to improve future estimates of root exudate C fluxes.

## Introduction

Root exudation – the release of low-molecular weight organic carbon (C) compounds from living plant roots into soil – mediates plant-soil interactions in a variety of ways, including facilitating plant nutrient acquisition via chemical and biological mechanisms (*e.g.* Jones and Darrah 1994; Meier et al. 2017), altering soil microbial communities (*e.g.* Shi et al. 2011), and impacting soil carbon dynamics (*e.g.* Yin et al. 2014). Root exudates mediate plant nutrient acquisition via a suite of chemical and biological mechanisms, in the process influencing the cycling of soil nutrients including phosphorus (P) and nitrogen (N). For instance, organic acids can increase plant access to inorganic P via rhizosphere acidification and phosphate mineral dissolution (Gillespie and Pope 1991; Hoffland 1992) or by displacing phosphate ions bound to mineral sorption sites (Jones and Darrah 1994; Bolan et al. 1994). Root exudates can also regulate mineralization of organic N and P by interacting with soil microbial communities. Enhanced root exudation has been correlated with the activities of enzymes involved in microbial N mineralization (Phillips et al. 2011; Meier et al. 2017) and P mineralization (Spohn et al. 2013).

Additionally, root exudation represents an important C flux to the soil that has unique effects on soil C dynamics. This is because root exudates can exert rapid effects on “stable” soil C formation and loss (more rapid than leaf or root litter which must be decomposed) (Sokol et al. 2019a). For example, chemical binding of C-based exudates onto soil minerals can result in soil C formation (Jones et al. 2003; Sokol et al. 2019b), whereas microbial activation due to fresh exudate C inputs could result in soil C loss via the “priming effect,” in which fresh C inputs stimulate microbial respiration rates (Kuzyakov et al. 2000). Recent field and lab experiments have shown that the rate of root exudation dictates both the rates of soil C formation and soil C loss (Yin et al. 2014; Chari and Taylor 2022) and is thus a relevant parameter for estimating soil C stocks and fluxes. In addition to its role in belowground C dynamics, root exudation may also be important for regulating C uptake aboveground. For example, in the Duke Free-Air CO_2_ Enrichment (FACE) experiment, aboveground productivity was sustained under elevated CO_2_ in part due to enhanced exudation and accelerated N turnover (Phillips et al. 2011; Drake et al. 2011). In addition to elevated CO_2_, warming, drought, and N deposition have all been shown to influence root exudation rates over relatively short timescales (*e.g.*, Phillips et al. 2011; Calvo et al. 2019; Xiong et al. 2020). Due to its role in regulating C cycle dynamics and its potentially rapid response to global change, capturing the exudation rate accurately at the global scale is necessary for making C cycle projections.

Process-based C cycle models often incorporate root exudation indirectly as a fraction of net or gross primary productivity (NPP or GPP, respectively) into a dissolved or low-molecular weight C pool (*e.g.*, Abramoff et al. 2018; Tao et al. 2023). The magnitude of this flux can be estimated in different ways. A number of modeling studies have used experimental measurements to constrain the direct C flux from plants to dissolved C pools (Sulman et al. 2014; Wieder et al. 2018). At least one model (Fixation and Uptake of Nitrogen, or FUN) calculates the root exudate flux as a function of C supply and nitrogen (N) demand, allowing it to be coupled to soil C models (Brzostek et al. 2014; Sulman et al. 2017). While we have the ability to link these models to local site-specific exudation data, we currently have virtually no empirically derived ecosystem-scale estimates of exudate C fluxes with which to compare the output of C cycle models at large scales. Given the increasing recognition of the importance of exudates in the C cycle and the increasing effort to incorporate this flux into ecosystem models, it is critical that we establish an empirical benchmark for the exudate C flux.

One reason for the absence of an exudate C flux estimate is the challenge of making exudate C measurements *in situ*. Exudates are released from live fine roots belowground, rapidly assimilated by microbes, and occur at low concentrations relative to large soil C pools resulting in small signal to noise ratios. To maximize the accuracy of *in situ* root exudation measurements, researchers must isolate intact fine roots from soil and incubate them under conditions that simulate the soil environment. Since the 2010s, *in situ* collection of root exudates has most frequently been performed using the “cuvette method,” which involves the incubation of a live fine root system in an open chamber filled with a glass bead matrix and nutrient solution culture (Phillips et al. 2008). Due to the challenges of accurately capturing root exudate C, relatively few *in situ* literature measurements exist compared to other major C fluxes. Here, we use a combination of these literature-derived measurements along with our own measurements of *in situ* root exudation to estimate the global root exudate C flux and biome-level exudate fluxes. We provide a) estimates for root mass-specific exudation rates, b) estimates for area-specific exudation rates, c) estimates for exudation as a proportion of GPP, and d) an estimate of the global annual flux of root exudate C into soil. Given the paucity of exudation measurements that exist currently, our estimates will be useful for improving the accuracy of simulated exudation rates in artificial root exudate experiments, incorporating root exudation into process-based C models, and improving ecosystem C budgets.

## Methods

### Data collection

We collected data from 40 studies measuring root exudation *in situ* including 128 sets of measurements (Fig. 1). Thirty-three studies were derived from a literature review and 7 studies were original, unpublished measurements by the authors. The literature review was restricted to studies of *in situ* (*i.e.* we excluded hydroponic studies and greenhouse experiments) total organic C exudation measurements from mature trees, shrubs, or grasses (*i.e.,* we excluded seedlings) using the cuvette method (Phillips et al. 2008). We used the search terms “in situ” “root exudation” and “cuvette method” on Google Scholar and additionally searched citations of Phillips et al. (2008) (the original description of the cuvette method).

**Fig. 1.**
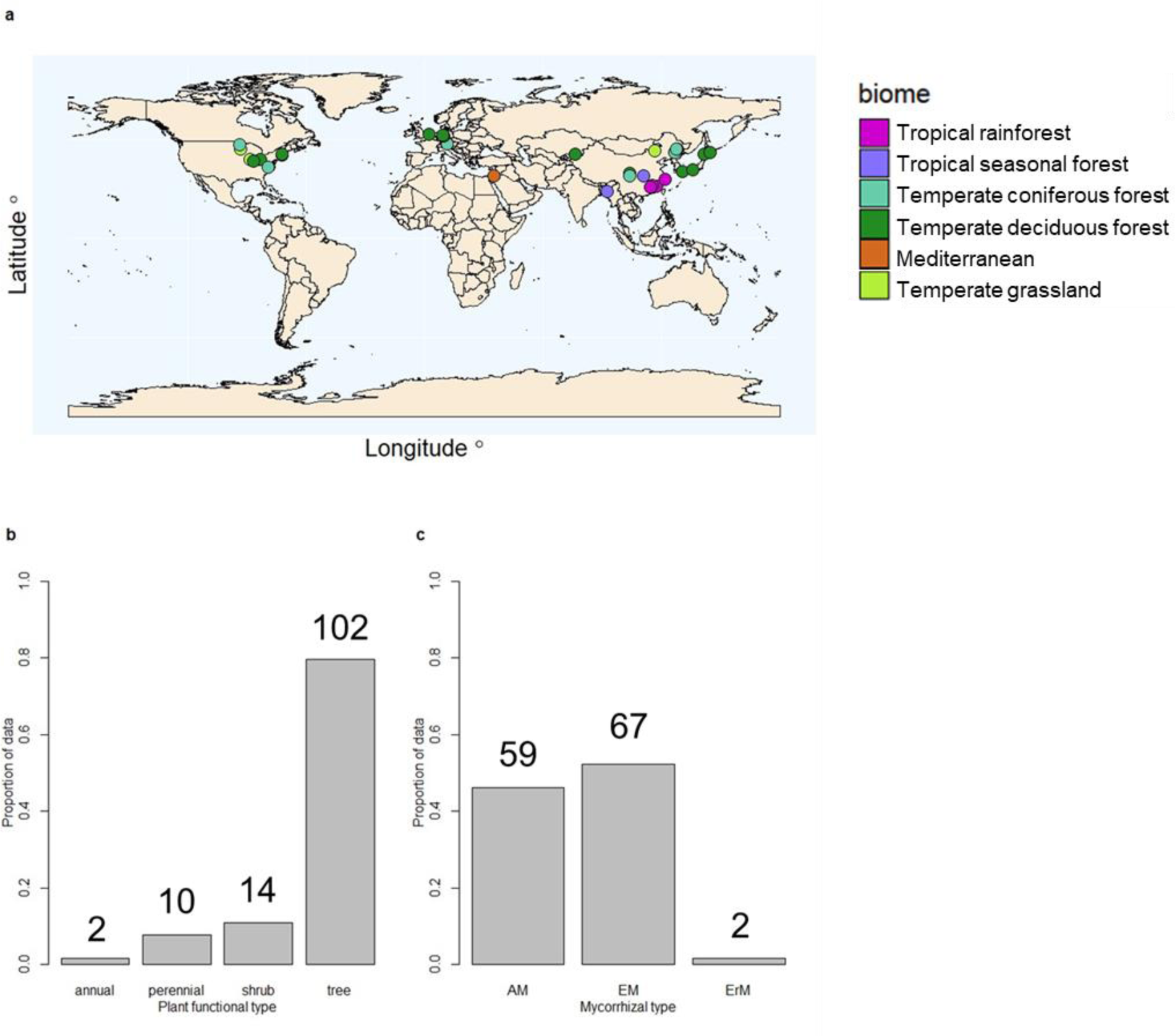
a) distribution of sites in the dataset on the globe colored by biome, b) proportional distribution of plant functional types in the dataset, c) proportional distribution of mycorrhizal types in the dataset. The numbers in b) and c) are the number of observations of each type.

We collected exudation rate data from each study in units of root exudate C per unit time per root biomass, root surface area, or root length. For each study, we determined the study-average exudation rate and the species-average exudation rate if multiple species were measured in a single study. When calculating the species-average rate, a small number of species with low sample sizes had a disproportionate influence on our calculations (*e.g.,* 4 species from the same study exceeded all other values in the dataset and exceeded the mean exudation rate by 4 s.d.), so we used the study-average exudation rate data for our scaling and analyses. Exudation rates are most commonly reported on a root biomass basis, which facilitates their inclusion into ecosystem C budgets. For studies reporting root exudation on a root surface area or root length basis, we converted these measurements to a root biomass basis using conversion factors derived from Jackson et al. (1997). Root exudation per unit root biomass per unit time (ug C g^-1^ h^-1^) is henceforth referred to as the specific exudation rate (SER).

### Scaling

We took a biome-level approach to scaling root-mass based exudation rates, which first required assigning each study in our SER dataset to one of 10 terrestrial biomes with fine root biomass data in Jackson et al. (1997) (Table 1) based on vegetation type and environmental characteristics. Of these 10 biomes, 6 are represented in the SER dataset. We then scaled the SER to a soil area basis using biome-level live fine root biomass (FRB) per soil surface area (kg m^−2^) estimates from Jackson et al. (1997). In this scaling, we made the assumption that *in situ* root exudation measurements are made from absorptive fine root biomass (AFRB). We estimated the proportion of total FRB that is AFRB for each biome (Table 1) from McCormack et al. (2015). To scale the root exudate carbon flux from an hourly basis to a yearly basis, we assumed exudation rates during the non-growing season are 76% of the growing season root exudation rate based on the studies in our data set that included growing season and non-growing season measurements (n = 10). We estimated the length of the growing season based on the biome (Table 1; Churkina et al. 2005; Piao et al. 2007). Thus, the root exudate C flux per soil surface area per year (henceforth *F_ex_* in units of kg C m^-2^ y^-1^) is calculated as follows for each data point:

**Table 1.** Parameters used in determining the global root exudate carbon flux. *scaled using cross-biome median SER, ^+^scaled using cross-biome median *F_ex_*, ^1^land surface area is included in woodland/shrubland. SER = specific exudation rate; FRB = fine root biomass; *P_AFRB_* = proportion absorptive fine root biomass; GS = growing season.

| Biome | Median<br>SER (ug C<br>g<sup>-1</sup> h<sup>-1</sup>) | FRB<br>(kg m<sup>-2</sup>) | $P_{AFRB}$ | GS<br>(d) | Land<br>surface<br>area<br>(10<sup>6</sup> km<sup>2</sup>) | Total flux<br>(PgC y<sup>-1</sup>) |
| --- | --- | --- | --- | --- | --- | --- |
| Mediterranean | 119.2 | 0.28 | 0.33 | 365 | NA<sup>1</sup> | NA<sup>1</sup> |
| Temperate<br>coniferous forest | 12.5 | 0.50 | 0.33 | 150 | 5.0 | 0.1 |
| Temperate<br>deciduous forest | 47.4 | 0.44 | 0.33 | 125 | 7.0 | 0.4 |
| Boreal forest | 67.4* | 0.23 | 0.33 | 130 | 12.0 | 0.5 |
| Temperate grassland | 69.9 | 0.95 | 0.81 | 100 | 9.0 | 3.5 |
| Savanna | 67.4* | 0.51 | 0.81 | 210 | 15.0 | 3.3 |
| Tropical deciduous<br>forest/seasonal<br>forest | 134.8 | 0.28 | 0.33 | 365 | 7.5 | 0.8 |
| Tropical evergreen<br>forest/rainforest | 180.5 | 0.33 | 0.33 | 365 | 17.0 | 2.9 |
| Woodland/shrubland | NA<sup>+</sup> | NA<sup>+</sup> | NA<sup>+</sup> | NA<sup>+</sup> | 8.5 | 0.7 |
| Desert | 67.4* | 0.13 | 0.81 | 200 | 18.0 | 1.0 |
| Tundra | 67.4* | 0.34 | 0.81 | 55 | 8.0 | 1.0 |
| Cultivated | NA<sup>+</sup> | NA<sup>+</sup> | NA<sup>+</sup> | NA<sup>+</sup> | 14.0 | 1.1 |
| **Total ($G_{ex}$)** |  |  |  |  |  | **15.2** |

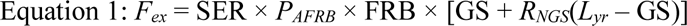

Where SER is the specific root exudation rate (expressed on a per-root-mass basis), *P_AFRB_* is the proportion of FRB in absorptive roots (as opposed to transport roots) (McCormack et al. 2015), GS is the length of the growing season, *R_NGS_* is the ratio of the exudation rate between non-growing season and growing season measurements, and *L_yr_* is the length of the year (Fig. 2).

**Fig. 2.**
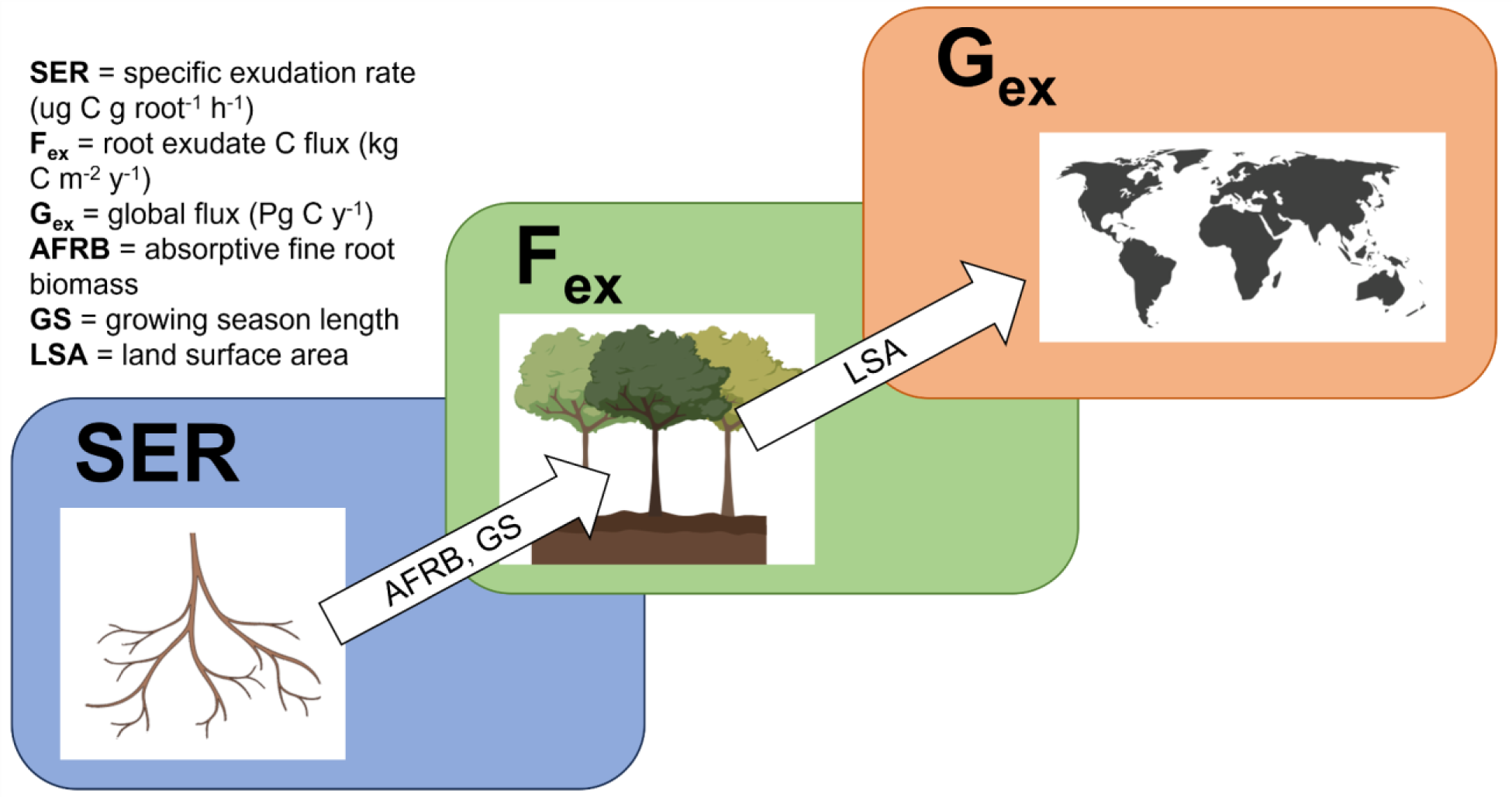
Conceptual figure illustrating the scaling process described in Methods. Specific root exudation rates (expressed on a per root mass basis) were scaled spatially using estimates of absorptive fine root biomass and temporally using estimates of growing season length to get exudate C flux estimates on a per-meter ground area basis (F_ex_). These estimates were then scaled to the entire biome using estimates of land surface area for each biome and all biomes were summed to get a global root exudate C flux estimate (G_ex_).

To determine per-biome average GPP we corrected per-biome NPP model estimates (Kicklighter et al. 1999) by an NPP:GPP factor of 0.46 (± 0.008 SE) (Collalti and Prentice 2019). We then divided *F_ex_* by the GPP of the appropriate biome to determine the proportion of GPP as root exudate C (*P_ex_*). Uncertainties for FRB, *P_AFRB_*, and NPP varied by biome and can be found in Table S2.

To determine the global exudate C flux, we scaled exudate C fluxes across land surface areas of each biome from Jackson et al. (1997). The 11 land area biomes differ slightly from the 10 FRB biomes presented in Jackson et al. (1997). Of the land area biomes, 5 overlap with FRB biomes from which we have SER measurements (temperate evergreen forest, temperate deciduous forest, temperate grassland, tropical seasonal forest, tropical rainforest), 4 overlap with FRB biomes without SER data (boreal forest, savanna, desert, tundra), and 2 do not overlap with FRB biomes (woodland/shrubland (which includes but does not overlap completely with the Mediterranean biome) and cultivated) (Table 1). For the 5 matching biomes, we scaled the median biome *F_ex_* by the global land surface area of the biome. For the 4 biomes that are not represented in our exudation dataset, we scaled the median SER of all biomes by the biome AFRB, biome GS, and biome global land surface area. For the two biomes that do not have FRB data, we scaled the median *F_ex_* of all biomes by that biome’s global land surface area. Two other scaling methods, which yielded similar results, are detailed in the supplement. We estimated the global root exudate flux (*G_ex_*) by summing the fluxes across the entire land surface area of each biome (Table 1, Fig. 2). To determine the proportion of global GPP, we divided *G_ex_* by a global GPP estimate from Badgley et al. 2019. All terms used in scaling to the global flux may be found in Table 1.

### Environmental data

We collected latitude and longitude information from each study in our dataset. We matched these coordinates to gridded global precipitation (Markus et al. 2022), temperature (Rohde and Hausfather 2020), and soil respiration (Stell et al. 2021) datasets to determine mean annual precipitation (MAP), mean annual temperature (MAT), and soil heterotrophic respiration (Rh) at each site. We analyzed relationships between SER and latitude, MAP, MAT, Rh, and mycorrhizal type (*e.g.*, plant associations with arbuscular vs ectomycorrhizal fungi) for both study-average and species-average data. We found that relationships between the species-average SER with latitude and MAT were due to several studies that sampled a high number of species at low replication (as explained above). We did not find these relationships with the study-average SER, which were less susceptible to anomalous values. As a result, only the study-average data are presented in the main text of this manuscript, and species-average relationships can be found in Fig. S1.

### Statistical analysis

We used ANOVA models to analyze differences in SER, *F_ex_*, and *P_ex_* between different biomes. We used Tukey’s Honest Significant Difference test to determine which biomes were different from each other. We used separate linear models to assess relationships between SER and each environmental predictor variable. All stats were done with the “stats” package. Significance was assessed at an alpha value of P = 0.05.

## Results

### Specific root exudation rate

The median SER across all biomes was 67.4 ug C g^-1^ h^-1^ (IQR = 17.8 – 107.6 ug C g^-1^ h^-1^). The mean SER is inflated due to non-normally distributed data; thus, we choose to report median SER here, which is likely closer to the true value (this is also true for all subsequent calculations). We did not find any statistically significant effects of biome type on SER (Fig. 3a). We found the species-average SER was greater in arbuscular mycorrhizal plant species than ectomycorrhizal plant species, but this difference was non-significant (*P* = 0.06) and disappeared when latitude was included as a covariate (*P* > 0.2). Study-average SER was greater in studies with only AM species than studies with only EM species, independent of latitude (*P* = 0.046 Fig. S2). We did not observe any effects of MAP, MAT, latitude, or soil heterotrophic respiration on the study-average SER (Fig. S1).

**Fig. 3.**
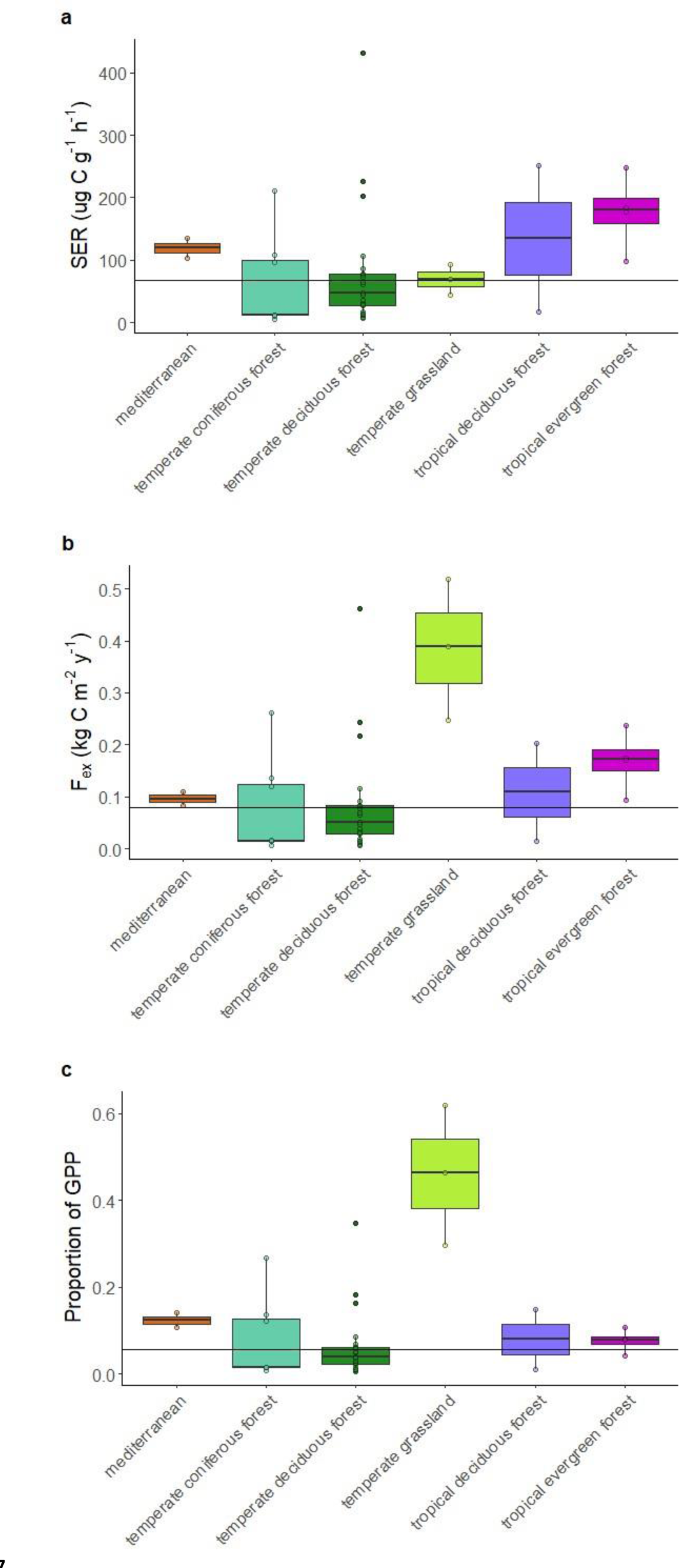
Exudation rates by biome. a) mass-specific exudation rate, b) soil area specific exudate C flux, c) proportion of GPP released as root exudates. For each biome, horizontal lines represent the median and boxes represent the inter-quartile range and whiskers represent 1.5 times the interquartile range. Colored dots represent individual data points (studies) used in our analyses and corresponding black dots represent outliers. Panel-wide black horizontal lines are cross-biome medians.

### Area-based root exudate C flux

We determined the root exudate C flux on a m^-2^ basis by scaling SER by AFRB. The median *F_ex_* was 0.079 kg C m^-2^ y^-1^ (IQR = 0.018 – 0.171 kg C m^-2^ y^-1^). *F_ex_* was significantly higher in temperate grasslands than temperate forests and Mediterranean biomes (*P* ≤ 0.040, Fig. 3b).

### Proportion of C allocated to exudates

We derived biome-level GPP estimates to determine the proportion of GPP as root exudates (*i.e., P_ex_* = *F_ex_*/GPP) within each biome. For the six biomes for which we could calculate *P_ex_*, the median *P_ex_* was 5.5% (IQR = 1.6% – 12.6%) and the median proportion of NPP was 12.0% (IQR = 3.5% – 27.4%). We found a significant effect of biome type on *P_ex_*, with temperate grasslands having higher *P_ex_* than all other biome types (*P* < 0.002, Fig. 3c).

We also compared our derived *F_ex_* to existing literature measurements of total belowground carbon allocation (TBCA) by mass balance for six field sites representing each biome in our SER dataset (Table 2). TBCA includes C allocated to root respiration, root production, rhizodeposition/exudation, and mycorrhizal allocation (Carol Adair et al. 2009). In theory, *F_ex_* should be some non-trivial proportion of TBCA, but should not exceed TBCA. Indeed, *F_ex_* ranged from 6% of TBCA (temperate coniferous forest) to 60% (temperate grassland) depending on the site.

**Table 2.** Comparison between mass-balance TBCA (total belowground carbon allocation) and root exudate C flux (*F_ex_*) (both in kg C m^-2^ y^-1^) in six field sites from biomes represented in this meta-analysis. Sources for TBCA are a) Almagro et al. (2010), b) Palmroth et al. (2006), c) Carol Adair et al. (2009), d) Litton et al. (2008), e) Raich et al. (2014). TBCA = total belowground carbon allocation; *F_ex_* = root exudate C flux per soil area.

| Site | Biome | TBCA (se) | $F_{ex}$ of biome (se) | Projected proportion of TBCA (se) |
| --- | --- | --- | --- | --- |
| Cehegín, Spain<sup>a</sup> | Mediterranean | 0.774 (0.035) | 0.096 (0.013) | 0.12 (0.02) |
| Duke FACE, NC, USA<sup>b</sup> | Temperate coniferous forest | 1.191 (0.083) | 0.073 (0.033) | 0.06 (0.03) |
| ORNL FACE, TN, USA<sup>b</sup> | Temperate deciduous forest | 0.782 (0.044) | 0.086 (0.023) | 0.11 (0.03) |
| BioCON, MN, USA<sup>c</sup> | Temperate grassland | 0.682 (0.019) | 0.385 (0.078) | 0.6 (0.1) |
| Kaupulehu dry forest preserve, HI, USA<sup>d</sup> | Tropical deciduous forest | 0.97 | 0.109 (0.095) | 0.11 (0.10) |
| La Selva, CR<sup>e</sup> | Tropical evergreen forest | 1.03 (0.20) | 0.169 (0.029) | 0.16 (0.04) |

### Global root exudate C flux

We scaled median biome-level data by land surface area measurements to determine the global root exudate C flux (Table 1). The median global root exudate C flux (*G_ex_*) was 15.2 Pg C y^-1^. We also scaled the first and third quartiles to obtain a range of 8.1 – 22.8 Pg C y^-1^. This flux represents 10.4% (range = 5.5% – 15.5%) of global annual GPP (147 Pg C y^-1^) (Badgley et al. 2019). While measurements in temperate forests account for 73% of the studies in our dataset, globally temperate forests only contribute 2.8% of the root exudate C flux. On the other hand, grasslands constitute only 3 measurements (< 8% of the dataset), but grasslands (temperate and savanna) represent 45% of the estimated global exudate C flux (Fig. 4).

**Fig. 4.**
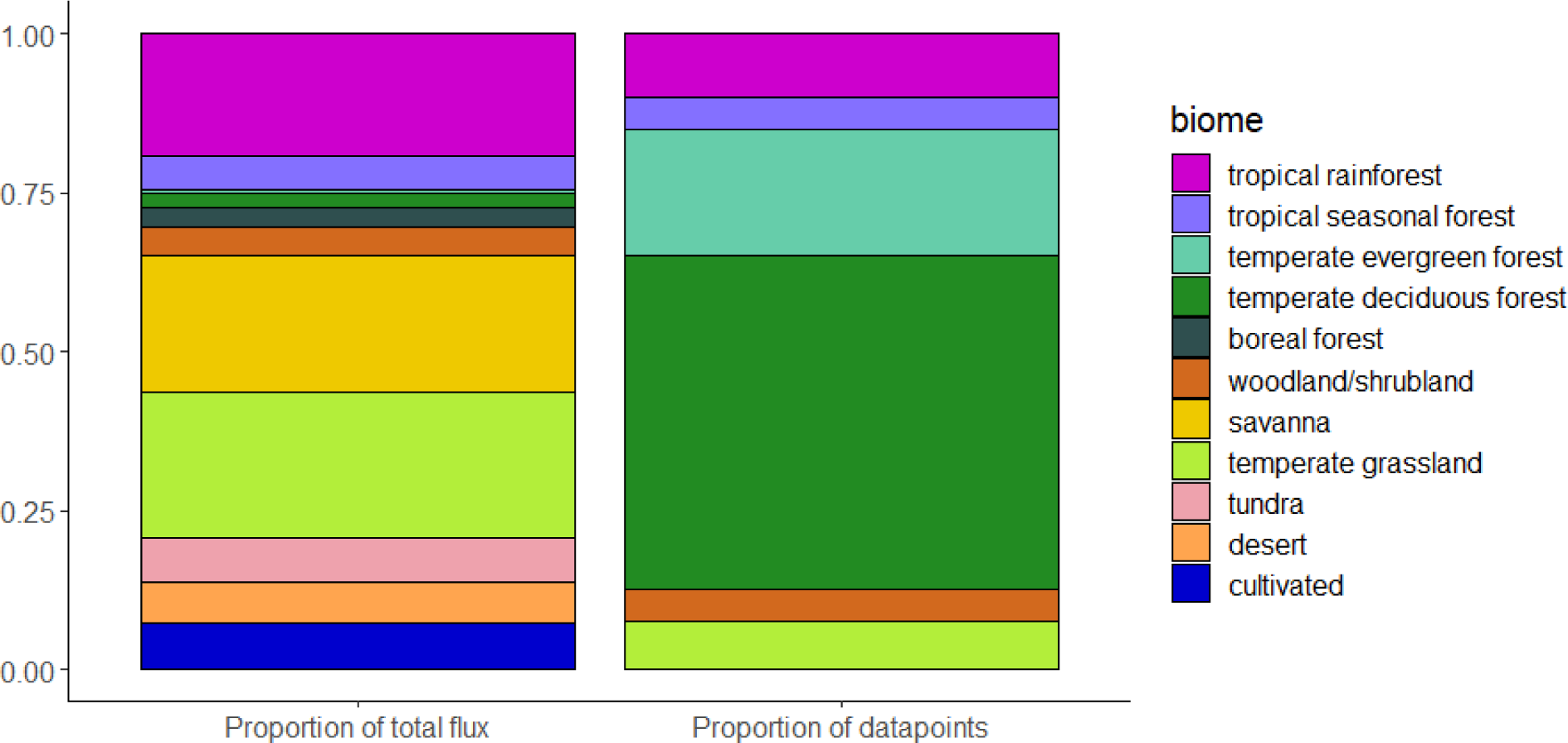
The proportion of the global root exudate flux from each biome (left) and the proportion of studies represented in our dataset from each biome (right). The left bar shows the proportional contribution of each biome to the global root exudate flux (Table 1). The right bar shows the proportion of studies in our dataset from each biome. The total number of data points for the right bar is 40.

### Sensitivity analysis

We varied each factor involved in our flux estimate independently to determine the sensitivity of the scaled estimate to each factor (Table S3). Per Equation 1, *F_ex_* is equally dependent on SER, FRB, and *P_AFRB_*. Of these three factors, *P_AFRB_* is the one that carries the most uncertainty, so an improved estimate of *P_AFRB_* is most likely to affect our flux estimates. Varying *P_AFRB_* by the range presented in McCormack et al. (2015) resulted in an uncertainty around the median *F_ex_* of over ±70%. *F_ex_* is also dependent, to a lesser degree, on GS and *R_NGS_*. However, varying these terms affected the median *F_ex_* by less than ±10%.

## Discussion

Our primary goal in this meta-analysis was to derive the most accurate and robust estimate of the annual global root exudate C flux possible given current data availability. Our analysis suggests a considerable global exudate flux of 15.2 PgC y^-1^ comprising 10.4% of global GPP. Before this study, the magnitude and importance of the root exudate C flux was largely unknown. At 10.4% of global annual GPP, the global root exudation flux (*G_ex_*) matches remarkably well with measurements from growth chamber experiments reporting 4 – 18% of photosynthetically fixed C released as root exudates (Barber and Martin 1976; Dror and Klein 2022). The amount of C released as exudates is likely to affect soil microbial activity, soil nutrient availability, and soil C dynamics, all of which affect feedbacks to plants and ecosystems at the global scale – and the results of this study suggest that this amount is considerable. Thus, we impress the importance of continued measurements of root exudation, studies which investigate the effects of root exudates on soil C dynamics, and incorporation of the root exudate C flux into models.

The global proportion of GPP that we estimate as root exudation (10.4%) differs from the median proportion of GPP in all the studies in our dataset (5.5%) – a difference that underscores the importance of grasslands to the global root exudate carbon flux. Grassland observations make up only a small proportion of our dataset (3 out of 40 studies), but together temperate grasslands and savannas represent 20% of the global land surface area (Jackson et al. 1997) and 45% of *G_ex_*. Grasslands contribute much more to *G_ex_* than their surface area would suggest because they have a much higher proportion of absorptive fine root biomass (*P_AFRB_*), where exudation primarily occurs, than forest ecosystems. The SER in grasslands is not significantly higher than other biomes, but because grasslands have both high FRB and *P_AFRB_*, the *F_ex_* is much higher in grasslands at 46% of GPP (compared to the overall median of 5.5%) (Fig. 3). Additionally, our TBCA comparison showed that exudation could be around 60% of TBCA at the grassland site, compared to 6-16% at other sites. While these numbers appear high, grasses invest most of their productivity belowground (Sun et al. 2021) and do not allocate C to build transport roots (McCormack et al. 2015), so grasses may be able to expend more C on absorptive root exudation. This is also consistent with stable isotope experiments which show most grassland belowground C export is associated with heavy fraction soil (where exudates accumulate) rather than light fraction soil (where root litter accumulates) (Fossum et al. 2022). However, we do caution that field-collected exudation data is extremely rare for grasslands (3 studies), so our estimates for this particularly important biome are based on relatively little current information (Fig 4).

Our estimate for the global exudate C flux (15.2 PgC y^-1^) is similar in magnitude to the estimate of 13.1 PgC y^-1^ allocated to mycorrhizal fungi derived by Hawkins et al. (2023). Hawkins’ results suggest that ectomycorrhizal plants allocate a greater proportion of their productivity to mycorrhizae than arbuscular mycorrhizal plants. Interestingly, our results provide some evidence that arbuscular mycorrhizal species may allocate more C to exudation than ectomycorrhizal species (Fig. S2), which suggests a potential trade-off between exudation and mycorrhizal C allocation. Arbuscular mycorrhizal associated plants may be more incentivized to allocate C to exudation as their mycorrhizae do not provide them with a robust organic nutrient acquisition mechanism (Read 1991; Phillips et al. 2013). However, we note that we did not find differences in exudation across arbuscular and ectomycorrhizal species in temperate forests where these species most commonly co-occur, so this difference may be better ascribed to an environmental or phylogenetic pattern. Finally, it’s important to note that exudation measured via the cuvette method (all of the studies in this meta-analysis) can also include fungal exudates (Kaiser et al. 2015), so these two flux estimates could partially overlap.

From a methods perspective, our results suggest that the cuvette method is robust for approximating root exudation rates. The agreement of our results with TBCA measured by mass balance suggests that the cuvette method captures the root exudate C flux to an accurate order of magnitude. Had the estimates of *F_ex_* exceeded TBCA, further refinements to the method may have been warranted. Nonetheless, some important considerations about field-collected exudate C data remain. Below, we discuss potential applications of the exudate fluxes we present here and suggest future approaches aimed at improving the quality of root exudate data for future flux estimates.

### Using the estimates from this paper

The estimates provided in this paper have a variety of potential applications for guiding future empirical research and constraining parameters for future modeling efforts.

#### Artificial root exudate (ARE) experiments

In ARE experiments, artificial exudate solutions are used to simulate the effects of root exudation on soil biological and physicochemical properties. One limitation of ARE experiments is that it is challenging to *a priori* simulate an accurate root exudation rate, so the responses observed in ARE experiments may not always be applicable in nature. Here, we provide estimates that researchers working in a variety of biomes can use to set the rate of root exudation in their experiments. For example, researchers could apply the *F_ex_* value presented here for their biome of interest and scale this value by the surface area of their incubation chamber and length of the experiment to know how much total artificial exudate C should be added.

#### Processed-based models

In process-based C cycle models, there is often a low-molecular weight carbon (LMWC) or dissolved organic carbon (DOC) pool (*e.g.,* Abramoff et al. 2018; Tao et al. 2023). Carbon can reach these pools either directly from plant input (*i.e.,* root exudation) or via microbial transformation of non-LMWC/DOC plant inputs. Since *P_ex_* is a function of GPP and models typically incorporate total C input to soil as some function of productivity, or GPP, modelers can use *P_ex_* to describe the proportion of plant input that moves directly to the LMWC/DOC pool without being microbially processed. We anticipate this flux may be larger than currently parameterized in processed-based models. For example, Wieder et al. (2018) incorporates exudation as 2% of NPP, or ∼1% of GPP compared to our median *P_ex_* estimates of 12% NPP and 5.5% GPP.

#### Ecosystem C fluxes

Ecosystem scientists and practitioners can use the estimates presented in this paper to constrain their estimates of root exudation without taking belowground measurements. For example, at field sites with remote monitoring of GPP from a flux tower, scientists can use our *P_ex_* estimates to determine an approximation of the C flux into soil as root exudates without on-site belowground field work. By comparing this estimate to TBCA as measured by mass balance (Palmroth et al. 2006; Carol Adair et al. 2009), researchers can approximate the proportion of C allocated belowground to exudation/rhizodeposition compared to fine root production.

### Data interpretation

We encourage readers to consider several caveats when interpreting these data. First, we stress that the SER was not different between biomes, and that differences in the exudation rate only emerged at the soil area level, due to high FRB and *P_AFRB_* in grasslands. Thus, our results do not show, for example, that grass roots exude more C than tree roots on a per-root basis. Rather, the enhanced *F_ex_* in grasslands is simply due to high absorptive fine root biomass. In fact, a key finding of our analysis is that there were no differences in SER across biome types, but this could simply be because a large majority of our data came from temperate forests. We call for more observations in non-temperate forest biomes to increase the statistical power of future analyses.

Additionally, we suggest that the estimates proposed in this paper are more likely to be overestimates than underestimates due to the way root exudation is measured. Commonly, root exudation rates are measured as the net release of C by a mass of fine root tissue into a cuvette over some incubation period (Phillips et al. 2008). Because exudates are measured from live fine roots, the cuvette incubation is inherently an open system and is thus prone to C contamination from outside sources, even though measurements are typically standardized with rootless “control” incubations. The consequence of this is that studies with a low sample size are particularly prone to the influence of a contaminated measurement. For this reason, we decided to focus on study-average rather than species-average data in this manuscript because presenting species-average data contained a high number of species with low sample sizes (*i.e.*, giving more weight to data with less replication). Biomes with a smaller number of studies in our dataset may be more likely to have a higher SER due to the same effect. This is also why we focused primarily on median rather than mean estimates.

Finally, we note that in scaling SER to *F_ex_* (and then to *P_ex_* and *G_ex_* subsequently) we relied on estimates of several other parameters including FRB, *P_AFRB_*, GS, and *P_NGS_*. Of these parameters, FRB was well constrained by Jackson et al. 1997 and GS and *P_NGS_* had relatively smaller effects on *F_ex_* (Table S3). *P_AFRB_* estimates, on the other hand, are the most uncertain (McCormack et al. 2015) and also carry equal weight as SER and FRB in determining *F_ex_*. Thus, we suggest our estimate is susceptible to change as *P_AFRB_* estimates are honed and encourage researchers to make *P_AFRB_* measurements when collecting root trait data.

### Future measurements

We urge researchers to continue taking *in situ* measurements of root exudation. Below, we outline several areas scientists can target in the future.

#### Improve measurement quality

Several steps can be taken to improve the quality of SER measurements. First, we strongly encourage researchers to prepare blank incubations and filter their samples when using the cuvette method. As a quality control method, researchers can look for a correlation between the total root exudate C and the root mass or root surface area. If there is no relationship, and specifically if low mass roots are generating high amounts of exudates, this is an indication that C contamination could be exceeding acceptable levels. C contamination could come from many sources, including both the environment and materials used in exudate collection. We urge researchers to be rigorous in checking for potential contamination before publishing measurements.

#### Collect exudates in under-represented biomes

Measurements of root exudation in temperate forests are over-represented in our dataset. From our current dataset, we are unable to determine if there are biome-level differences in the SER, largely due to substantial differences in data availability between biomes (Fig. 3a, Fig. 4). We encourage researchers to target biomes that have an overrepresentation in the global root exudate flux relative to an underrepresentation in the proportion of the dataset (Fig. 4). These include grasslands, tropical rainforests, agroecosystems, and all biomes in the global south. Improving data from these biomes should be the highest priority for constraining future global exudate C flux estimates. Agroecosystems merit special consideration as agricultural practices such as fertilization could have unique effects on exudation not observed in natural ecosystems.

#### Collect exudates seasonally, including outside the growing season

Exudation measurements are rarely taken outside of the plants’ dominant growing season, but these measurements are vital to determining the exudate C flux on annual timescales. Additionally, plants in different biomes exhibit reduced C assimilation during the non-growing season for different reasons. Since the plant response to the non-growing season is different between biomes where the non-growing season is driven by low precipitation vs. those driven by cold temperatures, unique estimates for the non-growing season rate in all biomes will be important for constraining the global exudate C flux.

#### Consider the effects of global change

Global environmental change affects numerous plant and soil processes, and a number of experimental measurements suggest drivers such as warming, drought, or elevated CO_2_ will affect the root exudation rate as well (*e.g.,* Phillips et al. 2011; Xiong et al. 2020; Brunn et al. 2022). If the estimates made in this paper are applied to climate change scenarios or experiments, estimates of root exudation would scale with GPP responses to climate change (*e.g.,* increase under eCO_2_ or decrease under drought). However, experiments suggest global change effects on exudation rates are not conserved in this manner – eCO_2_ has been found to decrease SER on numerous occasions (Dong et al. 2021), and drought to increase it (Calvo et al. 2019). We encourage scientists to continue measuring SER responses to climate change, and specifically to do so *in situ* in large-scale global change experiments with multiple drivers of change to help constrain these estimates.

#### Consider exudation in the context of root traits

Plants exhibit a suite of root traits, many related to nutrient or water acquisition, that can vary based on their individual physiology or environment. Most measured root traits are morphological (*e.g.,* specific root length, root tissue density, root diameter) or related to growth (*e.g.,* fine root biomass, production, turnover) (*e.g.,* Kong et al. 2019; Chen et al. 2021). Because exudation can be a plant strategy for acquiring both inorganic and organic soil nutrients (*e.g.,* Jones and Darrah 1994; Meier et al. 2017), we suggest researchers measuring root exudation incorporate it as a root trait in analyses of the root economic spectrum. Plants may trade off belowground C investment to exudation as opposed to structural C investment in root system expansion depending on their environment. Establishing relationships between exudation and other root traits may improve our ability to predict exudation rates and fluxes.

### Conclusion

This study represents the first effort to estimate the root exudate C flux at the biome and global scales. Our results suggest that root exudation is a considerable C flux into soils (roughly 10.4% of global GPP), that grasslands represent a relatively high exudate C flux due to their high fine root biomass, and that root exudation rates do not vary strongly across latitude or global gradients of temperature, precipitation, or soil heterotrophic respiration. We found some evidence that mycorrhizal associations impact root-specific exudation rates but note that studies where arbuscular and ectomycorrhizal plants co-occur showed no differences in exudation rates. Importantly, our analyses also indicate that measurements of this flux are data poor outside of temperate forests. Thus, we call for more measurements of root exudation in grasslands, tropical rainforests, and agroecosystems. Given the magnitude of the exudate C flux that our analysis suggests, we call for its continued and increasingly detailed incorporation into ecosystem C budgets and process based models.

## Author contributions

NRC and BNT devised the study, with input from SJT and RPP. NRC and BNT synthesized the literature data, and NRC, TLB, BDH, SO, TK, MKR, and SU collected original measurements. NRC analyzed the data with input and advice from BNT, SJT, and RPP. NRC wrote the first draft of the manuscript and all authors made editorial contributions to the manuscript.

## Data availability

The data analyzed in this study is publicly available at the following web address: [public web address to be supplied following acceptance of manuscript]

