## Supplementary Info for "The global root exudate carbon flux"

**Supplementary methods.**

*Determination of global exudate C flux.* We explored multiple methods for determination of the global exudate C flux ($G_{ex}$). The differences in these methods rest in how to determine an exudate flux out of biomes for which we did not have observations of SER in our dataset (boreal forest, savanna, desert, tundra). In the main paper, we scaled the median SER of the whole dataset by biome AFRB and land surface area to determine the flux out of these biomes (see *Methods*). The results of two alternative scaling methods (below) are shown in Table S1.

Method 2. Scale median $F_{ex}$ of whole dataset by biome land surface area. Here, instead of scaling the median SER by the AFRB of each missing biome (to derive a unique $F_{ex}$ for each biome), we scaled the median $F_{ex}$ directly. Effectively, this method assumes the AFRB of the missing biomes is not different from the median AFRB of the whole dataset. However, the remaining biomes generally had higher AFRB than the dataset median (McCormack et al. 2015), so we chose not to use this method for the main text.

Method 3. Instead of scaling the median SER by the AFRB of each missing biome, scale the SER of the "closest biome" to the missing biome (temperate coniferous forest for boreal forest, temperate grassland for savanna, desert, and tundra). We chose not to use this method in the main text because the classification of "closest biome" was subjective. However, using Method 3 resulted in almost the same result because there were no significant differences in SER by biome.

**Supplementary results.**

| **Method** | **$G_{ex}$ (PgC y-1)** | **Proportion of global GPP** |
|---|---|---|
| **1** | **15.2** | **10.4** |
| 2 | 13.7 | 9.3 |
| 3 | 15.1 | 10.2 |

Table S1. Comparison of alternative methods for determining $G_{ex}$. Method 1 is used in the main text.

| **Parameter** | **Mediterranean** | **Temperate coniferous forest** | **Temperate deciduous forest** | **Temperate grassland** | **Tropical deciduous forest** | **Tropical evergreen forest** |
|---|---|---|---|---|---|---|
| **FRB(SE) (kg $m^{-2}$)** | 0.28(0.096) | 0.50(0.10) | 0.44(0.053) | 0.95(0.078) | 0.28(0.049) | 0.33(0.050) |
| **$P_{AFRB}$(modeled range)** | 0.33(0.10 – 0.60) | 0.33(0.10 – 0.60) | 0.33(0.10 – 0.60) | 0.81(0.50 – 0.90) | 0.33(0.10 – 0.60) | 0.33(0.10 – 0.90) |
| **NPP(IQR) (kg C $m^{-2}$ $y^{-1}$)** | 0.358(0.256 – 0.388) | 0.452(0.410 – 0.564) | 0.612(0.507 – 0.790) | 0.385(0.335 – 0.482) | 0.632(0.539 – 0.701) | 1.012(0.937 – 1.098) |

Table S2. Table of parameters with uncertainty values that vary by biome.

| Parameter varied | Parameter range | Median $F_{ex}$ (range) (kgC m$^{-2}$ y-1) | Median proportion of GPP (range) | Median $G_{ex}$ (range) (PgC y$^{-1}$) |
|---|---|---|---|---|
| **SER** | SE = 14.4 (derived from dataset) | 0.079 (0.064 – 0. 094) | 0.055 (0.044 – 0.067) | 15.2 (12.4 – 18.0) |
| **FRB** | SE (from Jackson et al. 1997, see Table S2) | 0.079 (0.065 – 0.089) | 0.055 (0.049 – 0.062) | 15.2 (12.4 – 17.9) |
| $\boldsymbol{P_{AFRB}}$ | Modeled range (from McCormack et al. 2015, see Table S2) | 0.079 (0.024 – 0.144) | 0.055 (0.017 – 0.101) | 15.2 (7.4 – 21.5) |
| **GS** | ± 1 month | 0.079 (0.077 – 0.081) | 0.055 (0.054 – 0.057) | 15.2 (14.9 – 15.5) |
| $\boldsymbol{R_{NGS}}$ | SE = 0.11 (derived from dataset) | 0.079 (0.072 – 0.084) | 0.055 (0.051 – 0.060) | 15.2 (14.3 – 16.1) |
| **NPP** | IQR (from Kicklighter et al. 1999, see Table S2) | | 0.055 (0.043 – 0.067) | |
| **NPP:GPP** | SE = 0.008 (from Collalti and Prentice 2019) | | 0.055 (0.054 – 0.056) | |

Table S3. Sensitivity analysis of parameters involved in scaling calculations. Each parameter was varied by the range depicted while all others were held constant.

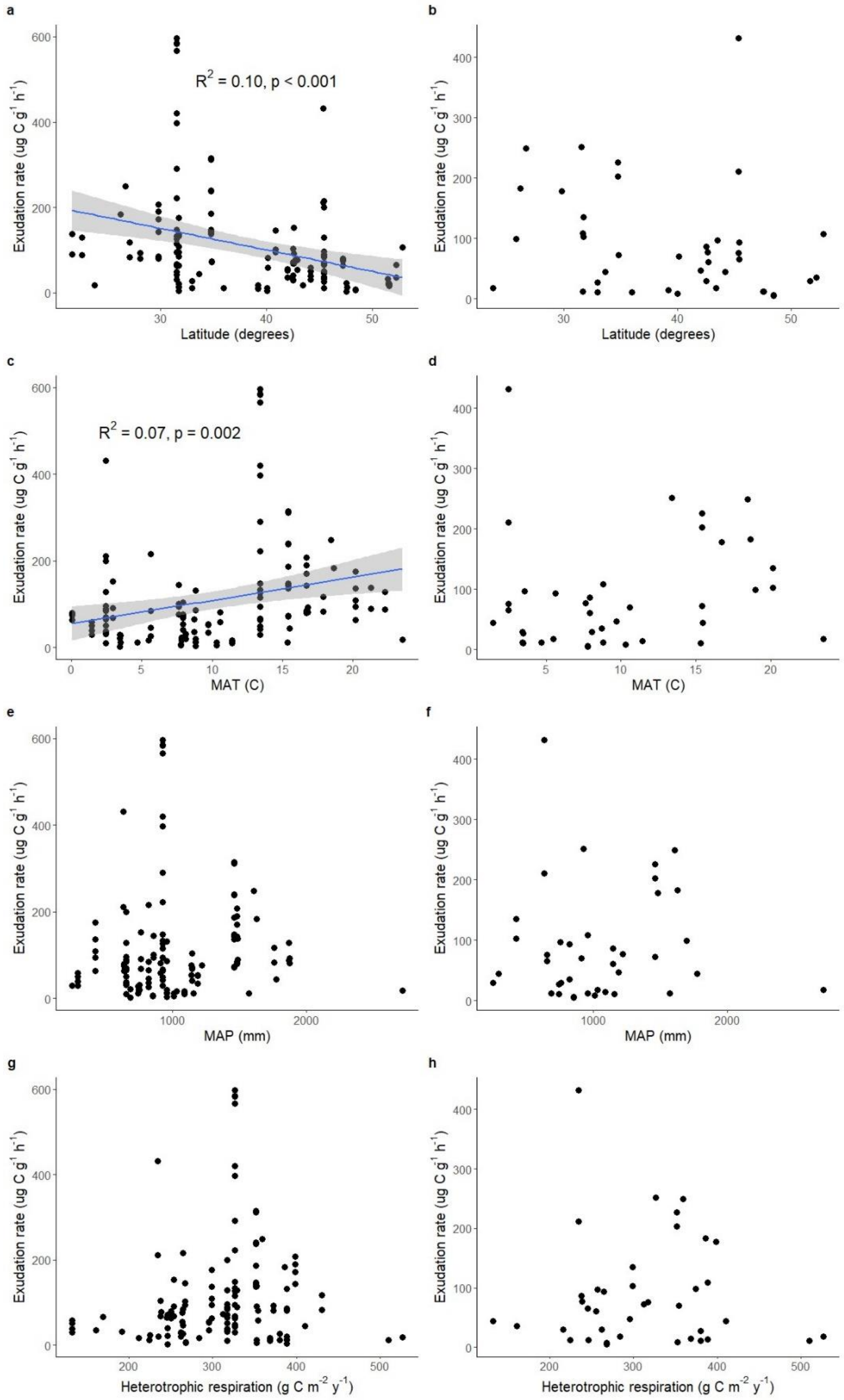


Fig. S1. Relationships between latitude (a,b), mean annual temperature (c,d), mean annual precipitation (e,f), heterotrophic respiration (g,h) and the exudation rate. Species-average relationships are shown in the left-hand column, study-average relationships in the right-hand column. The trendlines in (a) and (c) are from linear regression models.

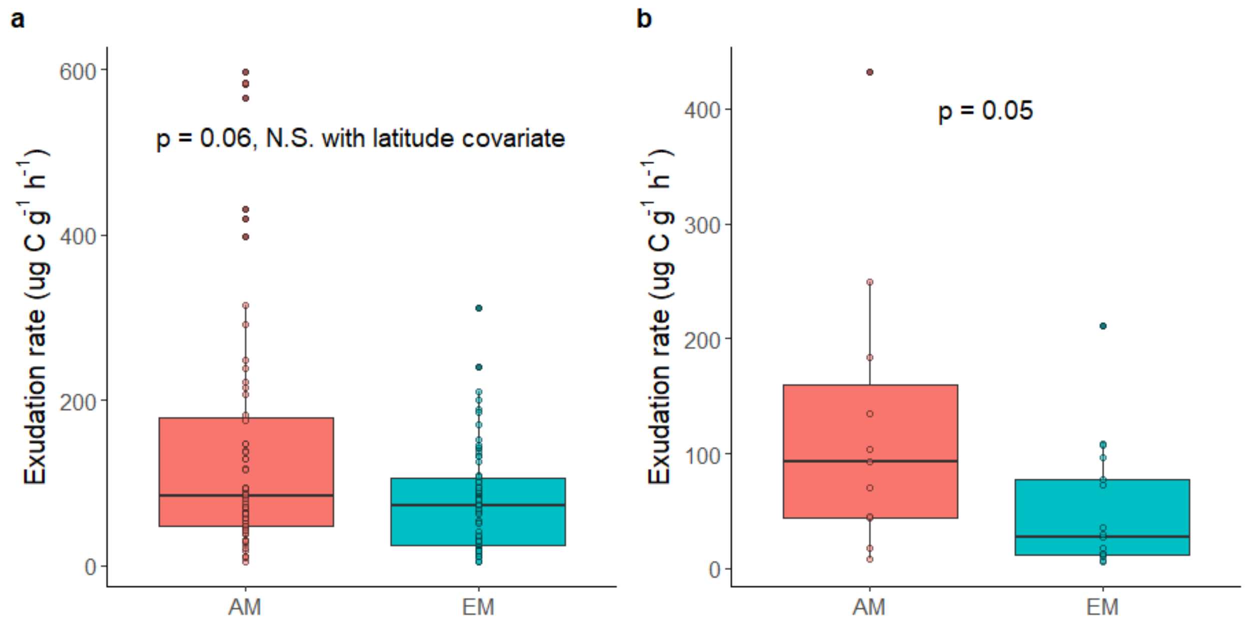


Fig. S2. Exudation rate by mycorrhizal type for species-average (a) and study-average (b) data. For (b), only single-mycorrhizal type studies are included. Horizontal lines represent the median and boxes represent the inter-quartile range and whiskers represent 1.5 times the interquartile range. Colored dots represent individual data points (studies) used in our analyses and corresponding filled black dots represent outliers.
